# Deciphering the tissue-specific regulatory role of intronless genes across cancers

**DOI:** 10.1101/2022.02.21.481319

**Authors:** Katia Aviña-Padilla, José Antonio Ramírez-Rafael, Gabriel Emilio Herrera-Oropeza, Guillermo Romero, Octavio Zambada-Moreno, Ishaan Gupta, Maribel Hernández-Rosales

## Abstract

**I**ntronless genes (IGs) or single-exon genes lacking an intron are found across most Eukaryotes. Notably, IGs display a higher transcriptional fidelity as they are not regulated through alternative splicing, suggesting better predictability biomarkers and easier regulation as targets for therapy. *Cancer* is a complex disease that relies on progressive uncontrolled cell division linked with multiple dysfunctional biological processes. Tumor heterogeneity remains the most challenging feature in cancer diagnosis and treatment. Given the clinical relevance of IGs, we aim to identify their unique expression profiles and interactome, that may act as functional signatures across eight different cancers. We identified 940 protein-coding IGs in the human genome, of which about 35% were differentially expressed across the analyzed cancer datasets. Specifically, ∼78% of differentially expressed IGs were undergoing transcriptional reprogramming with elevated expression in tumor cells. Remarkably, in all the studied tumors, a highly conserved induction of a group of deacetylase-histones located in a region of chromosome 6 enriched in nucleosome and chromatin condensation processes. This study highlights that differentially expressed human intronless genes across cancer types are prevalent in epigenetic regulatory roles participating in specific PPI networks for ESCA, GBM, and LUAD tumors. We determine that IGs play a key role in the tumor phenotype at transcriptional and post-transcriptional levels, with important mechanisms such as interactomics rewiring.

## 1. Introduction

Most eukaryotic gene structures contain exons interrupted by non-coding introns that are removed by RNA splicing to generate the mature mRNA. Although the most prevalent class of genes in the human genome contain Multiple exon genes (MEGs), about 5% of the genes are intronless (IGs) or single-exon (SEGs) that lack introns. Due to the absence of introns and associated post-transcriptional splicing, IGs entail a higher transcriptional fidelity suggesting a potential role as clinical biomarkers and drug targets that deserve careful consideration in diseases such as cancer [1-3]. Several previous studies have identified the role of IGs in cancer [2, 4-7]. For example the *RPRM* gene increased cell proliferation and tumor suppression activity in gastric cancer [7]; *CLDN8 gene* is associated with colorectal carcinoma and renal cell tumors [6]; while *ARLTS1* is upregulated in melanoma; *PURA & TAL2* are upregulated in leukemia [2] and protein kinase *CK2α* gene is up-regulated in all human cancers [4];

A remarkable instance of IGs acting in a clinical role is *SOX11*, a member of *SOXC* (SRY-related HMG-box) gene family of transcription factors involved in embryonic development and tissue remodeling by participating in cell fate determination [8]. *SOX11* has been associated with tumorigenesis, with aberrant nuclear protein expression in Mantle Cell Lymphoma (MCL) patients [9-12]. This TF is not expressed in normal lymphoid cells or other mature B cell lymphomas (except Burkitt lymphoma), but it is highly expressed in conventional MCL, including the cyclin D1-MCL subtype [5]. Hence, *SOX11* represents a widely used marker in the differential diagnosis of MCL and other types of small B-cell neoplasias in the clinical hemato-oncology practice [13-14].

However, to date, a comparative analysis integrating transcriptomic and interactomics profiles of IGs across diverse types of cancer is missing. Hence, this work aims to identify and characterize their expression, functional role, and interactomics profiles across different selected cancers.

## 2. Results

In this study we use RNA-sequencing data from 3880 tumor samples belonging to 8 cancer types from the cancer genome atlas or TCGA (**Supplementary Table 1**). We selected the four most prevalent cancers, namely Breast Invasive Carcinoma (BRCA), Colon Adenocarcinoma (COAD), Lung Adenocarcinoma (LUAD), Prostate Adenocarcinoma (PRAD); along with the four most aggressive cancers with high intrinsic heterogeneity, namely Bladder Urothelial Carcinoma (BLCA), Esophageal Carcinoma (ESCA), Glioblastoma Multiforme (GBM), and Kidney Renal Clear Cell Carcinoma (KIRC). We found that 338 out of the 940 genes identified as IG-encoded proteins are undergoing differential regulation in tumors compared to the normal tissue. GBM had the most number of DE-IGs at 168, followed by KIRC at 130, and both PRAD and COAD at 116. While in BRCA 104, LUAD 87, BLCA 97 and ESCA 86 were determined.

### 2.1 IGs tend to have a more induced gene expression pattern when compared to MEGs

In order to characterize the distinct biological behavior of IGs, we studied the overall gene expression patterns of IGs compared to those of the multi-exonic genes (MEGs) in all the tumors. Notably, a greater percentage of upregulated genes are found among DE-IGs than DE-MEGs in all the different types of cancers analyzed.

For instance, when comparing both populations of genes, we identified statistical significance for an over-representation of the up-regulated IGs over the MEGs group among DEGs in PRAD (*p-value*=2.8108 e-07); ESCA (*p-value*=7.0858 e-05); LUAD (*p-value*=0.0015), and BRCA (*p-value*=0.0276) cancers. Moreover, we aimed to compare if the upregulation levels in means of expression ranges are higher in IGs than in MEGs transcripts. Our analysis revealed that in almost all the cancers under study, except in lung adenocarcinoma, IGs tend to express in higher ranges than MEGs, as shown in **Supplementary Figure 1**. Further, we found a high proportion of IGs encodes for histone proteins involved in chromatin structure and other proteins essential to the regulation of development, growth, and proliferation (**Supplemental Figure 2**).

### 2.2 Upregulated IGs across cancer types encode for highly conserved HDAC deacetylate histones involved in negative gene regulation

To dig insight into the role of the prevalent upregulation mechanism identified for the DE-IGs, we aimed to characterize the groups of induced IGs across the analyzed cancer types. 338 out of 940 (∼35%) IG encoded proteins in the human genome, are differentially expressed across the eight analyzed tumor types. In a high number of them, 222 (∼78%), upregulation is conserved in two or more cancers, which suggests they are undergoing transcriptional reprogramming with higher rates of upregulated levels of their mRNAs as an outcome. Moreover, this upregulation mechanism is highly shared among the different cancers **(Figure 1)**. Most of the genes upregulated in BLCA and ESCA are shared with the other tumors, while tumors with a significant but less shared upregulation are GBM, PRAD, and KIRC, which could be expected given their remarkable heterogeneity (**Figure 1a**).

**Figure 1.**
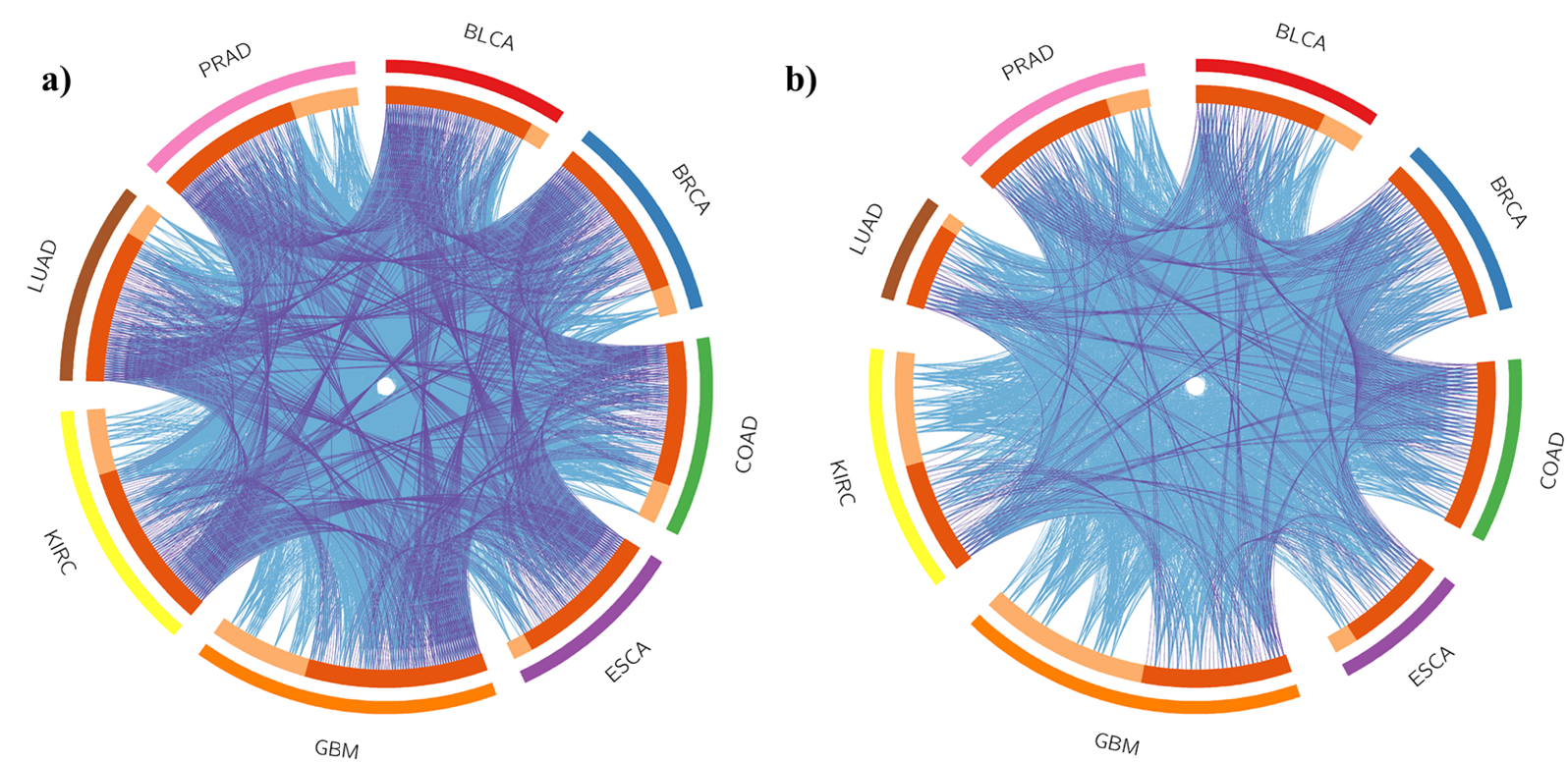
Deregulated IGs across the analyzed cancer genomes. The Circos plot depicts how deregulated genes for each cancer overlap. On the outside shell, each arc represents a cancer genome: BLCA in red, BRCA in blue, COAD in green, ESCA in purple, GBM in orange, KIRC in yellow, LUAD in brown, and PRAD in pink. On the inside shell, the dark orange color represents the proportion of DE genes shared with other cancers, while the light orange color represents the proportion of genes that are uniquely deregulated in a cancer. Purple arcs represent shared DE genes, while blue arcs represent shared functional enrichment terms among cancers. The greater the number of purple arcs and the longer the dark orange area imply a greater overlap among the deregulated IGs across cancers. (a) shows the upregulated genes, while (b) the downregulated.

To delve insights into the conservation of the activation of this negative gene expression mechanism in the cancer genomes, we identify the most conserved upregulated histones in all the diseases. We identified a group of 13 histone-related genes whose expression is highly conserved among the cancers (**Supplementary Figure 3)**. Moreover, this group of upregulated genes is statistically enriched in a proximal region (26.1968-27.8927 position in Mbps) in chromosome 6 in the human genome (**Supplementary Figure 4)**.

Haspin was found to be the only DE-IG shared among all the tumor genomes, highlighting its relevance in tumorigenesis. This gene is known to be required for histone H3 phosphorylation, necessary for chromosomal passenger complex accumulation at centromeres of mitotic cells [15-16]. In addition to its chromosomal association, it is associated with centrosomes and spindles during mitosis. The overexpression of this gene is related to a delayed progression through early mitosis [15].

Functional enrichment analysis was carried out for the deregulated IGs in all the studied diseases (**Supplementary Table 2**). GBM is the disease with the most DE-IGs and with the most diverse functional roles for the induced genes. The GBM-specific enriched terms are pathways of neurodegeneration, cell-cell adhesion, and gland development. The cancer-specific enriched terms chaperone-mediated protein folding, and regulation of neuron apoptotic processes were found for esophagus and colon cancers, respectively. Further, the Reactome pathways *R-HSA-321481* deacetylases histones (HDACs), and *R-HSA-3214858* RMTs methylate histone arginines were also enriched in IGs while GO:0006335 term: DNA replication-dependent chromatin assembly was also enriched suggesting an essential role in cancer biology.

### 2.3 IG downregulation is conserved in breast and colon cancers and is involved in signaling and cell-specific functions

When comparing the downregulated IGs and their enriched functional terms among the diseases, it was pointed out that the breast and colon cancers share all the repressed IGs with their functional pattern. Downregulated genes in those cancers are involved in the regulation of cellular localization, sensory organ development, regulation of membrane potential, adipogenesis, epithelial to mesenchymal transition in colorectal cancer, growth, actin filament base process, positive regulation of cell death, class A/1 (rhodopsin-like receptors), and the regulation of secretion by cell biological processes, (**Supplementary Table 2**). Notably, the most shared downregulated pathway among cancer genomes is the regulation of anatomical structural size, shared among BRCA, PRAD, GBM, LUAD, and BLCA tumors. Additionally, unique roles are found in lung and prostate cancers. They have downregulated genes involved in the vitamin D receptor pathway, and wounding response respectively.

### 2.4 Cancer-specific differentially expressed IGs

To study the specificity in the regulation of human IGs, we quantified the number of cancer-specific DE-IGs (**Figure 2**). We also built a bipartite network to identify cancer-specific and shared deregulated IGs, where upper nodes represent DE-IGs, and bottom nodes represent cancer types, we link an IG to a cancer if it was found to be deregulated in that cancer (**Supplementary Figure 2**). Among the 338 DE-IGs, we identified that ∼35% were specific for a cancer type **(Supplementary Table 3)**. GBM followed by KIRC and PRAD with 35, 18, and 16 DE-IGs respectively displayed the highest specificity for IG expression.

**Figure 2.**
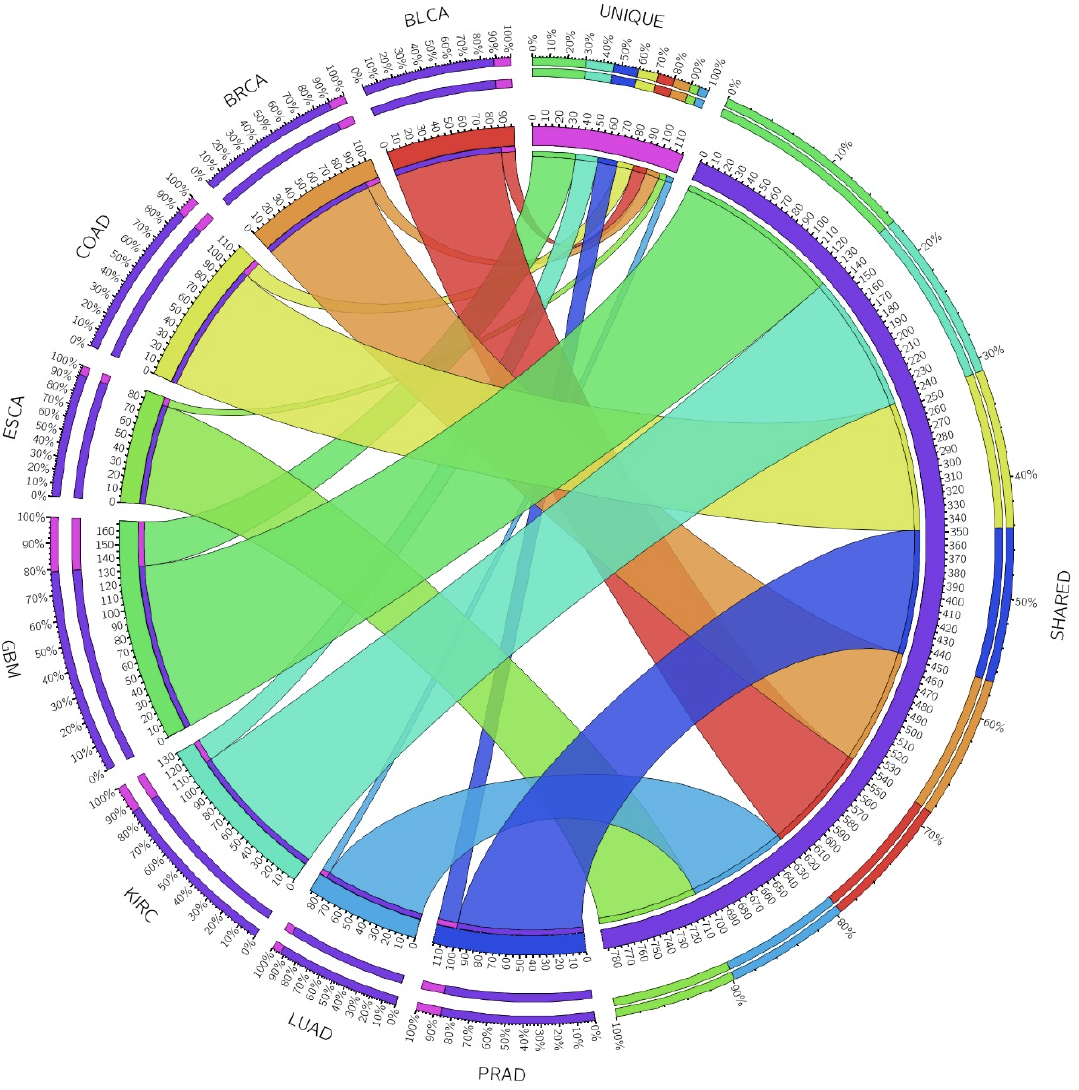
Shared and unique differentially expressed IGs across cancers. Circa plot depicts the number of shared DE-IGs across the 8 types of tumors under study. In the left hemisphere, the total number of DE-IGs in each cancer is represented by an arc: BLCA (red), BRCA (orange), COAD (yellow), ESCA (light green), GBM (green), KIRC (turquoise), LUAD (light blue), and PRAD (dark blue). The unique (fuchsia) and shared (purple) groups are depicted in the right hemisphere. The inner arcs represent the distribution of the DE-IGs in each cancer, while the outside arcs represent the intersection of both hemispheres on a scale of 0-100%. Data was obtained from the TCGA database.

#### 2.4.1 GBM

Among the specific DE-IGs found in GBM, an evident first group includes mainly cellular proliferation and cytoskeleton-related genes. An enrichment of microtubule-based movement, transport along the microtubule, cytoskeleton-dependent intracellular transport, and microtubule-based transport is identified (FDR= 0.0132). Moreover, the second group of specific deregulated genes is related to tumor suppression, a negative regulatory mechanism of cell growth control, which usually inhibits tumor development. We identified GBM specific deregulated genes with crucial involvement in the *p53* pathway, such as *KLLN, INSM2* and *SFN*.

#### 2.4.2 KIRC

The cancer-specific deregulated IGs in KIRC include the tissue-specific *KAAG1* gene (kidney-associated antigen 1) along with transcription factors *POU*5, *TAL*, and *bHLH*. Moreover, transmembrane proteins such as the protocadherin beta 10 (PCDH10), related to the Wnt signaling, were also upregulated. *PCDH10* has been classified as a tumor suppressor in multiple cancers. It induces cell cycle retardation and increases apoptosis by regulating the *p53/p21/Rb* axis and *Bcl-2* expression [17].

#### 2.4.3 PRAD

IGs specifically downregulated in PRAD include *RAB6D* (member RAS oncogene) and *MARS2* (methionyl-TRNA Synthetase 2) related to amino acid metabolism, while *PBOV1*, a 135 amino acid protein with a transmembrane domain was highly upregulated (logFC=3.1699). This gene is mapped to 6q23-q24, a region associated with loss of androgen dependence in prostate tumors [18]. Other upregulated IGs include transcription factors such as *FOXB2* (Forkhead Box B2), and *ATOH1* a *bHLH*, which is associated with goblet cell carcinoid and Merkel cell carcinoma diseases (Verhaegen et al 2017). Moreover, *FRAT2*, an essential positive regulator of the Wnt signaling and *LENG9*, a member of the conserved cluster of receptors of the leukocyte-receptor complex (LRC) were also upregulated [19].

#### 2.4.4 BRCA

Few cancer-specific differentially expressed IGs that code for TFs were identified for BRCA, while specific gene expression is more abundant for transmembrane proteins in this cancer, such is the case of a group of glycoproteins and transmembrane receptors involved in signal transduction. For instance, upregulation of *PIGM* (phosphatidylinositol glycan anchor biosynthesis class M), *GP5* (glycoprotein V platelet,) *CALML5* (calmodulin-like 5), coiled-coil domain-containing proteins *CCDC96*, and *CCDC87*, are identified specifically for this cancer.

#### 2.4.5 LUAD

Regarding the specific differential expression found for lung tumors, the following TFs are significantly upregulated: *TAF1L*, a transcription factor that is related to chronic inflammatory diseases by regulating apoptotic pathways including regulation of *TP53* activity, and *FOXE1*, a forkhead TF previously related to thyroid cancer [20]. Interestingly, specific transmembrane signaling receptors are upregulated. *MAP10* is a microtubule-associated protein and *THBD* thrombomodulin endothelial-specific type I membrane receptor that binds thrombin. Among *THBD* related pathways are collagen formation and formation of fibrin clot, as well as calcium ion binding and cell surface interactions at the vascular wall.

#### 2.4.7 BLCA

In BLCA, only ZXDB (Zinc Finger X-Linked Duplicated B) TF is identified as specifically deregulated. Transcripts for signal transduction proteins are specifically upregulated for *OR52E8 (*Signaling by GPCR), *LRRC10B* (Leucine-Rich Repeat Containing 10B), and *MAS1L*, a proto-oncogene which is a peptide ligand-binding receptor linked to sensor neurons for pain stimuli detection.

#### 2.4.6 ESCA & COAD

No specific TFs are specifically differentially expressed for colon and esophageal cancer. Gastrointestinal adenocarcinomas of the tubular gastrointestinal tract, including esophagus, stomach, colon, and rectum, share a spectrum of genomic features, including TF-guided genetic regulation [21]. In keeping with this, we found that transmembrane proteins related to GPCR signaling pathways are specifically undergoing differential expression for both cancers; for instance, in colon, *GPR25* (G protein-coupled receptor 25), *CCDC85B* (coiled-coil domain containing 85B), *CCDC184* (coiled-coil domain containing 184), *SPRR1A* (small proline-rich protein 1A), *PROB1* (proline-rich essential protein 1). Meanwhile, for esophagus tumors, *OR51B4* related to signaling by GPCRs, and *LDHAL6B* involved in glucose metabolism and respiratory electron transport were identified.

Remarkably, when comparing IG’s deregulation across the eight cancer types, we identified that ESCA and LUAD shared 93% of the DE-IGs, while BRCA, BLCA, COAD, PRAD, and KIRC shared more than 86% at least with another cancer. Finally, and presumably given its *“multiforme”* nature, GBM is the cancer type identified with a significant but lesser percentage of shared DE-IGs (∼79%). Notwithstanding, our results show that GBM represents the tumor where the IGs have more differential expression, specificity, and concerted functional roles. This could be explained due to their gene expression tissue-specific relevance in the brain.

### 2.5 Proteins encoded by cancer-specific deregulated IGs interact with distinct groups of proteins in PPI networks

Gene expression is a phenomenon where coupled biochemical interactions take place to transcribe mRNA for protein production. There is a high regulation in the balance of this event. Hence differences in protein composition, production, or abundance are a consequence of disruptions in cell phenotypes. The differential expression pattern of the mRNA is a key in determining a cell state at the molecular and physiological level [22]. It has been shown that genes involved in *“similar diseases”* depict shared protein-protein interactions (PPI), and higher expression profiling similarity [23].

To determine if the DE-IGs play a crucial role in inhibiting or exacerbating biological reactions at a physical level, we obtained the Protein-Protein Interactions (PPIs) with highest confidence where DE-IGs for each cancer are involved. Our results show that a considerable proportion of proteins encoded by DE-IGs in each cancer interacts with specific groups of proteins, due to the low percentage of DE-IGs that are shared between two or more cancers (**Supplementary Figure 5**). In this work, we will consider that an interaction is shared between cancers, if a DE-IG in one cancer is also deregulated in another cancer, and the interaction of the IG-encoded protein with another protein is reported by String, otherwise, we classify that interaction as cancer-specific.

For instance, 100% of the interactions found in LUAD are cancer-specific, followed by the high specificity of interactions found in ESCA (99.3865%) with only a unique shared PPI of centromere complex with BLCA (*HIST1H2BJ, CENPA*). GBM has a similar pattern with 95.8333% of unique interactions and only 19 shared ones. In BLCA, 59.16% of the analyzed PPI are specific to this cancer type. PRAD is a condition that possesses 53.1339% cancer-specific interactions; and COAD has 42.1455%. For BRCA, 30.62% of its interactions are only found in this cancer, the lowest fraction of unique interactions. KIRC has the second-lowest with only 31.55% of unique interactions.

If two cancers share PPIs related to the cancer-specific DE-IGs, there could be an underlying affected process characteristic for such diseases. To determine those specific processes, protein complexes involved in shared interactions were examined, finding that most of the identified proteins belong to families of core histones. Overall, the analyzed PPIs indicate a very distinct pattern of interactomics for each cancer. The tumors that share the greatest proportion of PPIs are breast and colon, having 145 common interactions representing 44.61 % of the total interactions identified for such diseases. The second higher similarity found is for prostate and kidney tumors, sharing 227 interactions (28.64 % of all). The rest of the tumors share at most 21.32 % interactions (**Supplementary Figure 5, Supplementary Table 4**).

These cancer-specific interactions were analyzed deeper to delve into their functional role in each cancer (**Figure 3**). As it could be expected due to their high specificity at the PPI level, this network approach shows specific clusters well defined for glioblastoma (**Fig 3b**) and lung (**Fig 3d**). In less-defined clusters, esophageal and bladder cancer can be observed in panels **e)** and **f)**. In contrast, in the case of prostate, bladder, and kidney tumors, the interactions are linked by common DE-IGs in a cluster. On the other hand, we identified that the communities in the network are defined primarily by the type of proteins. Functional enrichment shows specific biological processes intrinsic to the physical interactions implied for the genes in each cancer (Supplementary Figure 6). For instance, proteins involved in BLCA-specific PPIs are conducting mainly chromatin organization and DNA repair reactions, while in colon tumors, proteins play a concerted role in the regulation of transcriptional processes. In keeping with this, in esophageal tissue, the interactors are involved in the regulation of different classes of non-coding RNAs. Meanwhile, in glioblastoma and lung tumors, proteins are specifically interacting for splicing activity, and transcription initiation processes, respectively. Kidney and prostate tumors show specific protein-protein interactions for DNA-replication and protein-protein complex formation, and DNA organization and packaging. In contrast, proteins with specific interactions in breast tumors, have an important activity linked to cellular growth and apoptotic processes.

**Figure 3.**
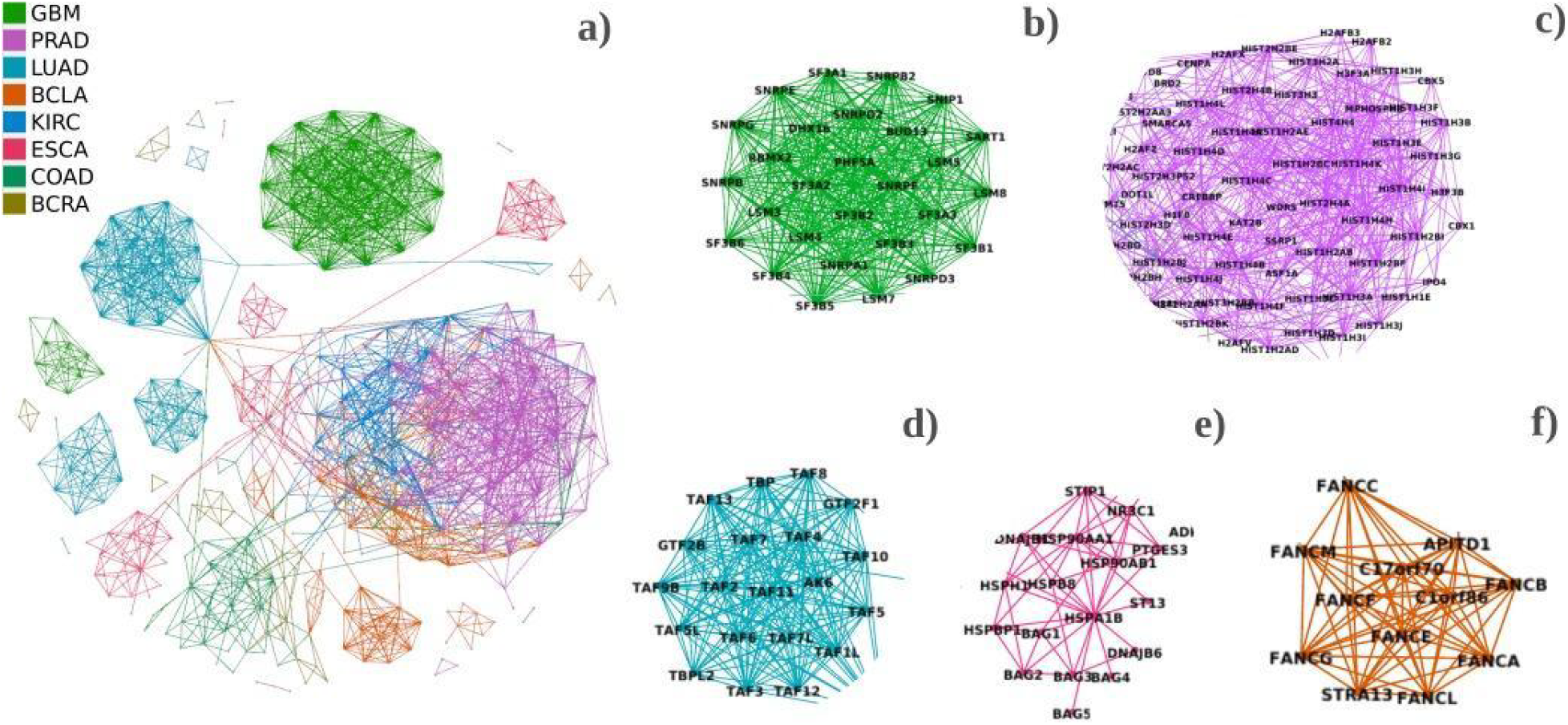
Protein-protein interaction network of DE-IGs and their 50 closest interactors across cancers. **a)** Unique and shared PPI across cancer are shown, each cancer has a specific color for its interactions. Close-ups of the biggest and more isolated clusters in the network are presented: b) GBM, c) PRAD, d) LUAD, e) ESCA, and f) BLCA. Data was obtained from the String database.

Intrinsic of the PPI networks is the topological information capable of characterizing cancer proteins [24]. Surprisingly, our comparison of IGs with their physical interactors at a protein-protein level shows a tendency to lower betweenness centrality, lower degree, and a few interactions with other IGs. Therefore, even though IGs are not hubs in the network of interactions where they participate, changes in their regulatory role can cause a cascade of disruptions in interactions that might lead to malignancy.

Altogether, these results are the opposite of the patterns reported for cancer genes, which suggests that the behavior tendency of IG proteins is to interact with *oncogenic* genes [24].

For a closer approach, the nearest fifty interactors of the DE-IG encoded proteins in each cancer were analyzed. In **Figure 4** we can observe that cancer-specific DE-IG encoded proteins interact with specific groups of proteins in distinct cancers, suggesting that DE-IGs affect particular post-transcriptional processes.

**Figure 4.**
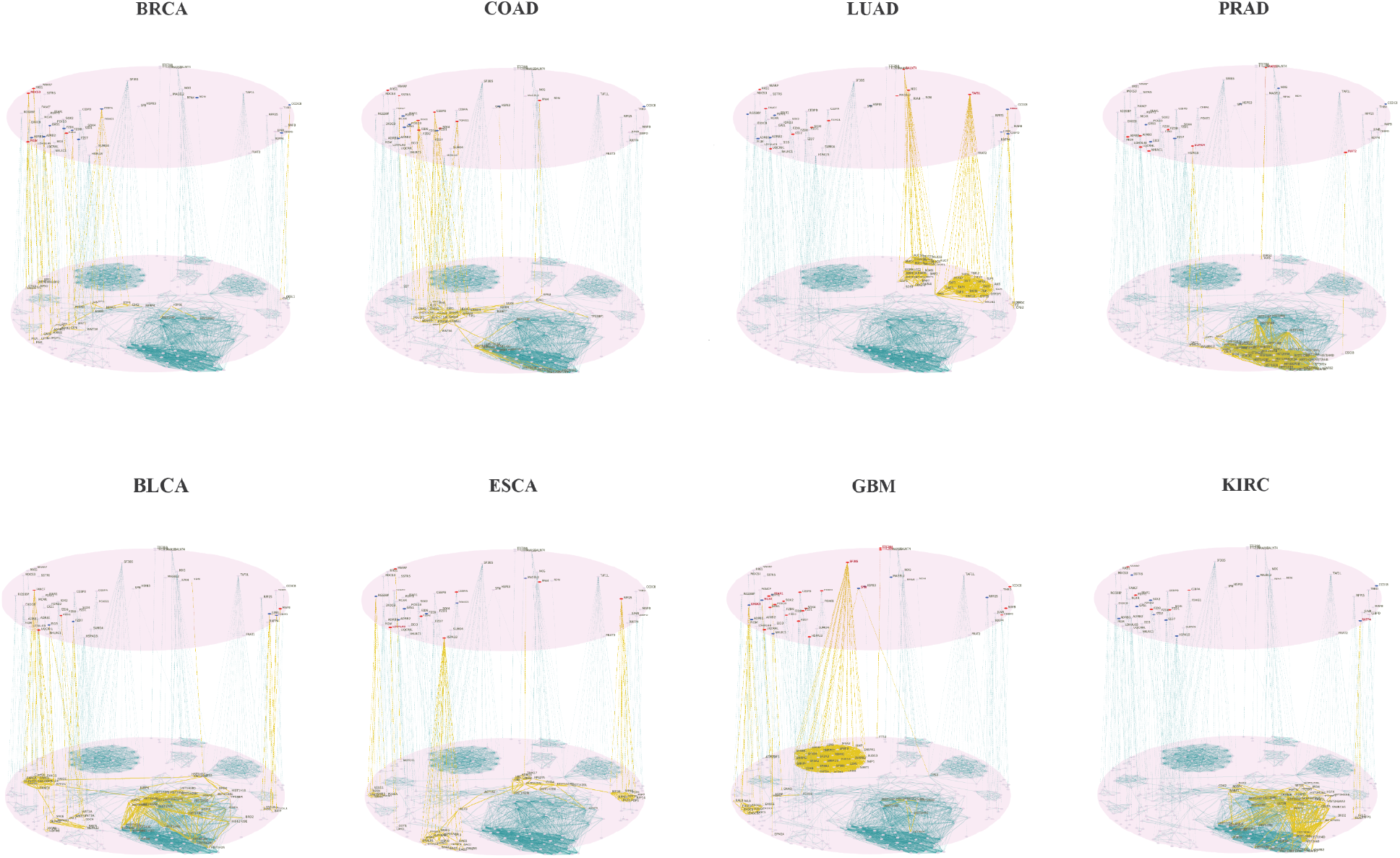
Multilayer networks of IG-encoded proteins and their cancer-specific interactors. Protein-protein interactions were found for deregulated IGs in each cancer. The cancer-specific deregulated IGs that encode for proteins interact with exclusive groups of proteins in each cancer. Upper layer shows deregulated IGs across cancer, highlighting the cancer-specific upregulated IGs in red,) and downregulated in blue. Links to the bottom layer connect IG-encoded proteins to proteins found in PPIs by String. Yellow links highlight interactions that can be affected by deregulation of IGs in a specific type of cancer.

The most important protein complexes for each type of cancer were determined for cancer-specific deregulated IGs and their interactors (**Supplementary Table 5**). In total, we detected 27 protein complexes where cancer-specific DE-IGs and their interactors are found. Colon cancer is the disease containing more DE IGs in the same complex: 9 out of 13 corresponding to the disassembly of the beta-catenin Wnt signaling pathway. Bladder DE IGs are interacting mainly with the proteins of the CD40 signaling pathway, and are involved in multi-multicellular organism processes. Likewise colon, breast cancer-specific IGs are related to wnt signaling pathway, but also to follicle-stimulating hormone signaling. Glioblastoma-specific proteins have a major participation in splicing, and vesicle transport. Kidney’s are more related to regulation of feeding behavior. Esophageal’s IGs are involved in protein folding, and tRNA and ncRNA processing. Lung cancer proteins participate in transcription preinitiation complex assembly, as well as in the BMP signaling pathway. Prostate cancer DE-IGs are found mainly related to chromatid sister cohesion and translation and postranslation processes. The protein complex related to Wnt signaling pathway is associated with a large group of deregulated genes among Bladder, Breast, Colon, Kidney and Glioblastoma, being a total of 17 IGs with disrupted expression.

### 2.6 DE-IGs participate in the genetic *“rewiring”* of cancer cells

Cancer cells undergo significant genetic “*rewiring”* as they acquire metastatic traits and adapt to survive in multiple environments with varying nutrient availability, oxygen concentrations, and extracellular signals. Therefore, to effectively treat metastatic cancer, it is important to understand the strategies that cancer cells adopted during the metastatic process. Finally, we focus on studying the “*rewiring”* between healthy and tumor samples using BRCA as a model (**Figure 5**). We used breast cancer since it is the only dataset that fits the criteria for a mutual information analysis (at least 100 samples for each condition are required). A network of DE-IGs and their co-expressed genes was built and analyzed for each condition. Our results show that the tumor co-expression network is composed of a total of 62,462 interactions among 15,347 genes, while the healthy network has 13,037 genes with 45,941 interactions. There are only 9,615 interactions shared between the two network topologies. All the differences among these two networks are potential *rewiring-caused* co-expression interactions that may be part of the mechanism that BRCA cells follow to achieve the characteristics of their phenotype. For instance, as seen in **Figure 5**, IGs like *RRS1, FNDC10, EPOP* and *NND* are highly affected by this *“rewiring”* behavior in cancer. According to differential expression and network analysis, we observe that histones play a key role in rewiring the co-expression mechanism from healthy to cancer tissue. Examples of this are H2AW and H2BC8, which are core histones that have only few healthy tissue-specific interactions that are almost lost during cancer development, which creates many new interactions with other genes.

**Figure 5.**
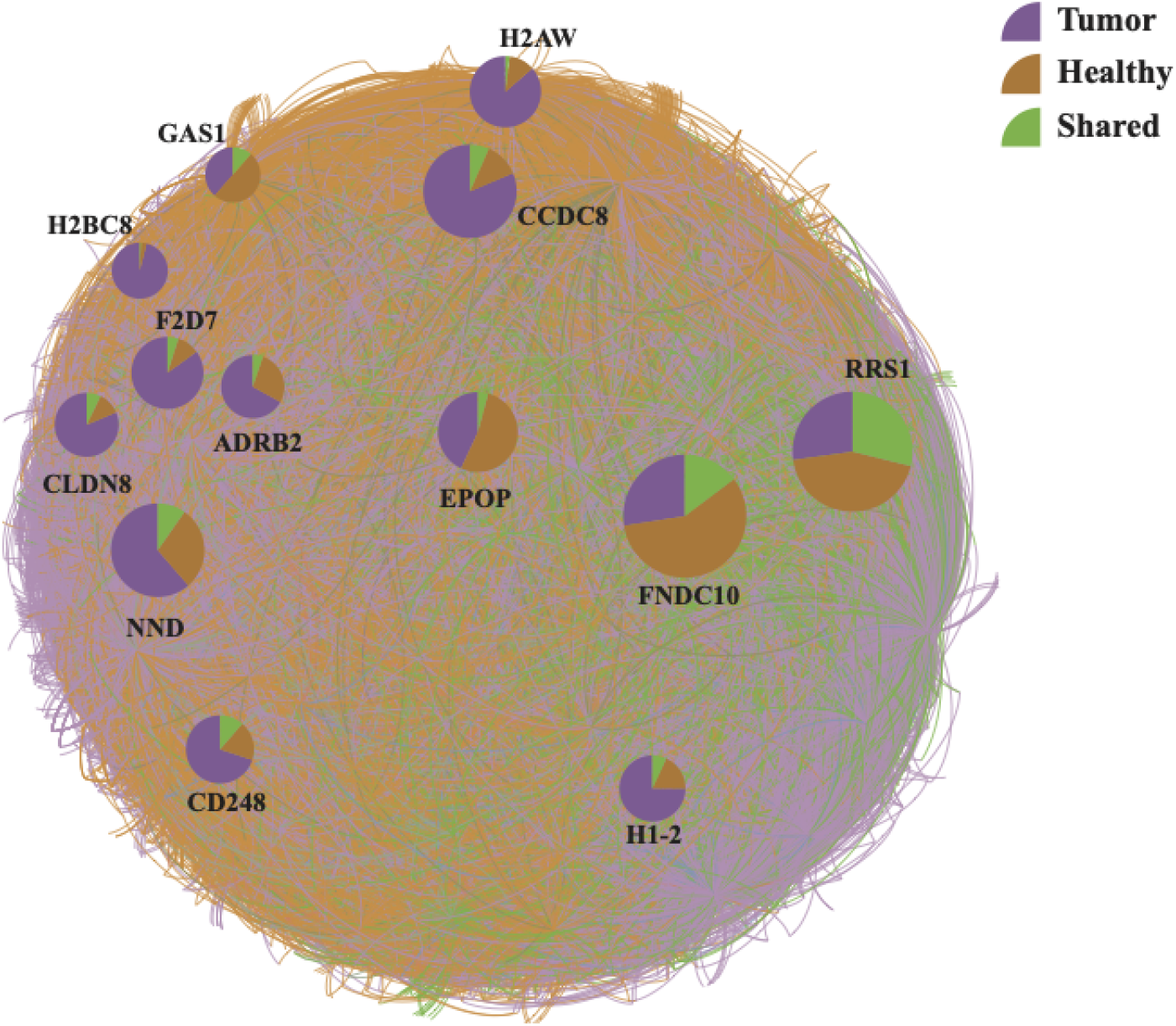
Co-expression network of healthy and tumoral breast tissue. In this network, the co-expression of DE-IGs found in BRCA in healthy tissue (depicted by brown links) and breast tumor tissue (purple links) is shown. Shared co-expression relations are depicted with green links. Pie charts for IG nodes of a higher degree are shown. Each of them indicates the percentage of each type of interaction: in purple tumor-specific interaction, in brown healthy-specific interaction, in green conserved interaction. We can see the effect of rewiring, by observing that a high percentage of interactions that are present in healthy tissue is not observed in tumor tissue. Moreover, many more new interactions emerge in the tumor tissue.

## 3. Discussion

Here, we present a comprehensive analysis of differentially expressed Intronless genes (IGs) across eight cancers using a standardized bioinformatics pipeline. We uncover signature changes in gene expression patterns of IGs such as enhanced expression changes when compared to multi-exon genes, unique and shared transcriptional signatures between different cancer types, and association with specific protein-protein interactions that might have potential effects in post-transcriptional processes.

Strikingly, our results show that the upregulated IGs across the eight different cancer types were related to negative regulation of gene expression (gene silencing) and negative regulation of cell differentiation. Further, we found a core set of chromatin-modifying genes such as *HASPIN* and Histone deacetylases (HDACs), located on chromosome 6, that were upregulated across all the analyzed cancers. Although the pattern of differentially expressed (DE) IGs was found to be not specifically shared across the different tumor types, highlighting their specificity modulating cancer signaling and proliferative pathways. For example beta-catenin Wnt signaling was regulated in kidney and prostate tumors while GPCR signaling critical to inflammation [25-27] was regulated in gastrointestinal esophageal and colon adenocarcinomas, and p53 pathways.

This biological behavior could be due to tumor’s cellular origin and heterogeneity [28], and tissue-specific expression [2, 8]. Due to their remarkable heterogeneity, glioblastoma, prostate, and kidney tumors were found with a less representative shared behavior in the upregulation of IGs. Notably, glioma tumors are the malignant microenvironment where IGs showed the most characteristic gene expression regulation profile with 35 unique DE-IGs, more than 50% greater than other cancers. Notwithstanding, our results show that GBM represents the tumor where the IGs have more differential expression, specificity, and concerted functional roles. This could be explained due to the “multiforme” nature of GBM that is driven by brain-specific gene expression as is further evidenced by least shared DE-IGs (∼79%) with other cancers.

In conclusion, IGs drive massive transcriptional rewiring, as observed in co-expression analysis, that may drive tumorigenesis and may act as novel therapeutic targets for gene therapy and in-silico drug design such as popular GPCRs or emerging HDAC inhibitors [29].

## 4. Materials and Methods

### 4.1 Data extraction and curation for IG, and MEG datasets

Data was extracted using Python scripts (https://github.com/GEmilioHO/intronless_genes), and the Application Programming Interface (API). *Homo sapiens* genome was assembled at a chromosome level and was accessed at the Ensembl REST API platform (http://rest.ensembl.org/ accessed using Python with the ensembl_rest package). The pipeline process was as follows: protein-coding genes with CDS identifiers for transcripts for all chromosomes were retrieved and classified into two datasets named “single-exon genes’’ (SEGs), and “multiple exon genes’’ (MEGs) depending on exon and transcript count. The SEG dataset was submitted to the Intron DB comparison (http://www.nextgenbioinformatics.org/IntronDB) to separate those with UTR introns and referred to as uiSEGs. The output of the pipeline was a third dataset containing only intronless genes (IGs). After data extraction, a manual curation step in the IG and uiSEG datasets was followed to discard incomplete annotated protein sequences and mitochondrial encoded proteins. The final IG dataset contained 940 protein-coding genes with only one exon and one transcript.

### 4.2 Bipartite network & quantification of shared and unique DE-IGs

The bipartite network was constructed with upper nodes representing IGs and bottom nodes representing cancer types. We place links connecting each IG to the cancer types where such a gene is differentially expressed (DE). Nodes corresponding to genes in the bipartite network were sorted by degree aimed to visually identify intronless genes cancer-specific deregulated, and those whose expression is disrupted in most of the diseases. A heatmap for shared IGs is computed by identifying the number of deregulated IGs in every pair of cancers, and dividing such quantity by the total of the disrupted genes in any of the compared diseases. This metric is known as the Jaccard similarity coefficient.

### 4.3 Data source and differential expression analysis across cancer

Currently, the (The Cancer Genome Atlas) TCGA possesses data for the study of 37 different tumor types. We found the primary tumor and adjacent normal tissue transcriptomic data for 18 types. Among those, we selected 8 types for the present study. Gene expression data from patients for BRCA, BLCA, COAD, ESCA, GBM, KIRC, LUAD and PRAD cancers was downloaded from the NIH website (https://portal.gdc.cancer.gov/) using the TCGAbiolinks R package [30] with the following restriction criteria: samples types primary tumor and solid tissue normal (control); results of RNAseq experimental strategy; and workflow type HTSeq-counts format. Differential expression analysis was carried out using the TCGAutilis R package [31], indicating which of the obtained samples correspond to tumors and which to control, and establishing a filtering threshold of FDR = 0.05 and logFC = 1 (absolute value) to consider a gene significantly differentially expressed.

### 4.4 Functional Enrichment Analysis of Differentially Expressed IGs

The functional enrichment was conducted using the over-representation analysis of the functional assignment (ORA). Genes with differential expression up to one log2-fold change values were considered as up-regulated with a *p-* and *q-value* set at 0.05 and 0.10, respectively. First, the functional enrichment of the 338 differentially expressed human IG proteins (up-regulated and downregulated separately) was performed using all human IGs proteins as a background *“universe”* (selecting input as species: *Homo sapiens*, universe 940 human IGs). The comparative functional enrichment analyses were performed using Metascape (https://metascape.org/) for the biological process category, including KEGG and Reactome pathways. To delve insight into the role of specific IGs in the affected biological processes ORA was assessed to determine category barplots. The Circos software [32] was employed for data analysis and visualization.

### 4.5 DE-IGs PPI network construction & protein complex identification

Network analysis and visualization were performed using python scripts (See repository) and the Gephi software. StringDB platform [33] was used to download physical and functional interactions data. The highest confidence scores (0.9) were filtered for this study, keeping the most probable interactions. Then, interactions were requested for the set of unique DE-IG of each cancer type, downloading relations between those genes, and also their interactors in the first shell (up to 50 interactors). All this data is assembled into a single network. Network metrics such as degree distribution, closeness, and betweenness centralities were computed using Python networkx library [34]. For protein complex detection, we assumed that highly clustered nodes in the PPI network correspond to protein complexes. Therefore, 27 protein complexes were predicted using the Louvain method to find communities implemented in Gephi.

### 4.6 BRCA network deconvolution

Co-expression networks were inferred using ARACNe-AP [45], a mutual information-based tool for gene regulatory network deconvolution. In this analysis, we used separated submatrices for healthy and tumor tissue and the list of every DE-IG in the BRCA dataset. For this study, a *p-value* of 1 × 10^−8^ was set up and 100 bootstraps were carried out. Then, they were consolidated into a single network, getting an inferred co-expression network for each condition.

## Acknowledgments

We would like to thank Roddy Jorquera and Carolina González for fruitful discussions. Special thanks to Fernando Flores for helping with designing and constructing the co-expression network. We are also thankful to Carlos González for visualization and technical support.

This research was funded by Conacyt Ciencia Basica Project 254206. K.A.P (CVU:227919), J.A.R.R (CVU: 1147711), and O.Z.M (CVU: 1147042) received financial support from CONACyT. K.A.P is a current holder of a fellowship from the Fulbright Comexus García-Robles foundation.

## Appendix

**Supplementary Figure 1.**
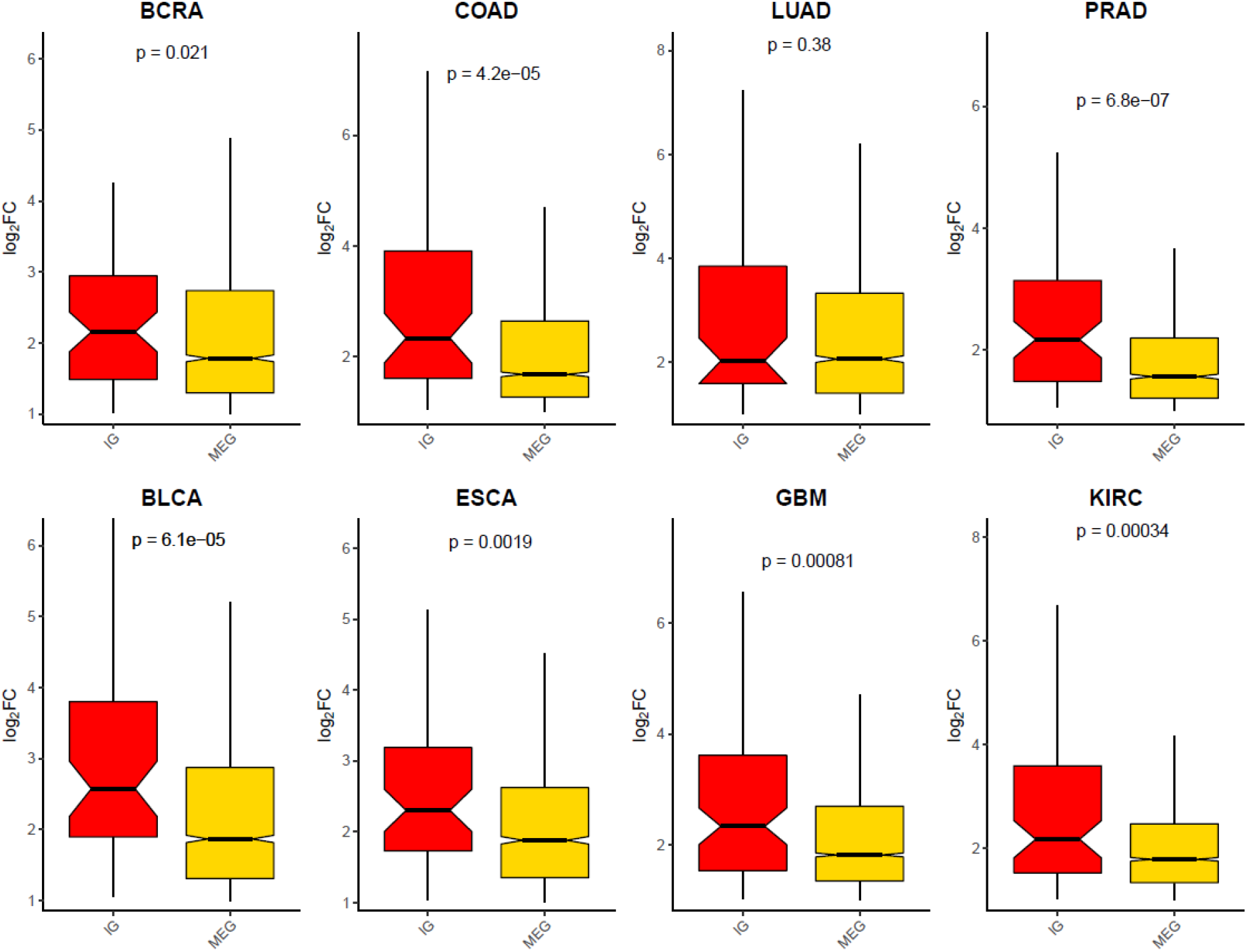
Upregulated gene expression ranges among cancers. A boxplot is shown for each cancer type, separated by gene type (IGs or MEGs); expression ranges are shown on the y-axis using log2fold change values. The boxes present a notch corresponding to the confidence interval for each population. In the upper part of the graph, the *p-value* for the Wilcoxon test is shown in order to indicate the statistical difference between both populations.

**Supplementary Figure 2.**
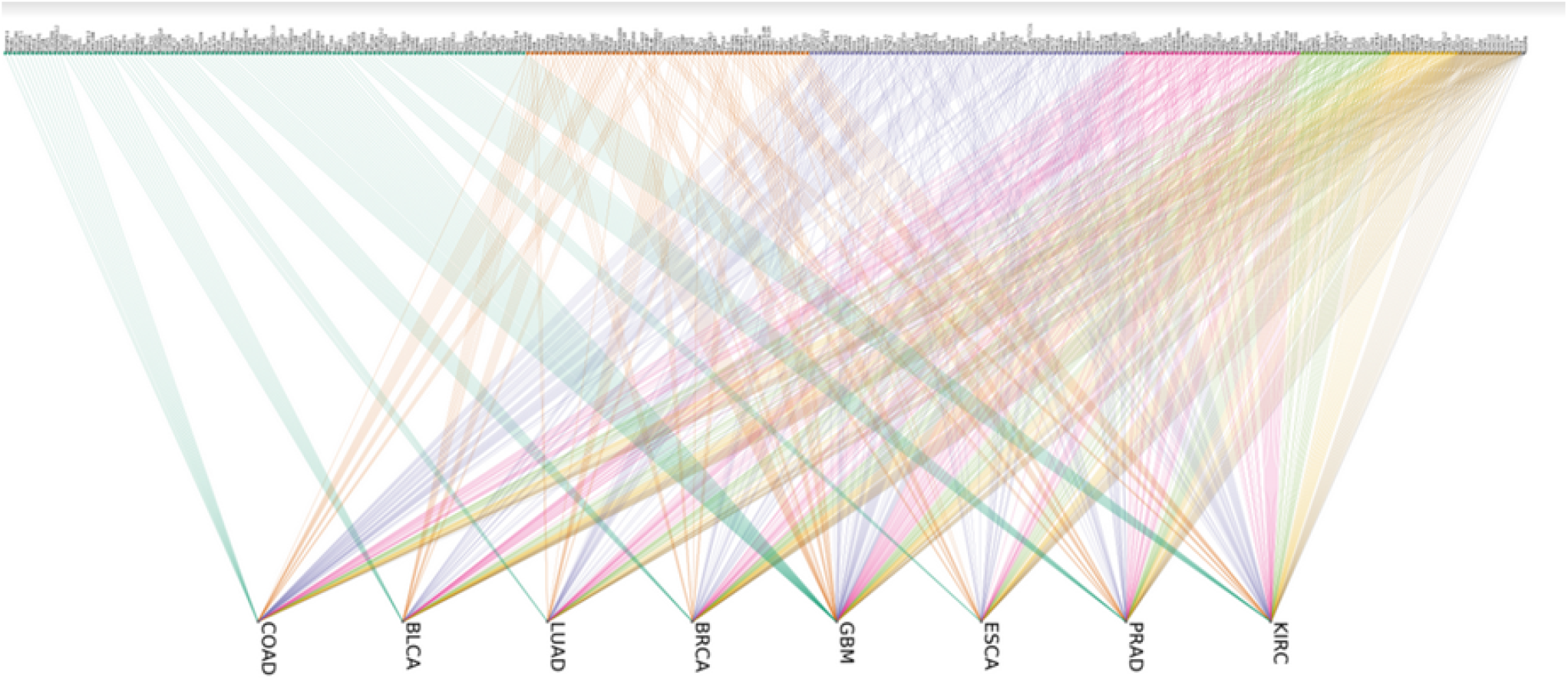
IG-cancer type bipartite network. Intronless genes are shown in the upper group of nodes, while cancers are located in the bottom group. A link joining an IG and a cancer type indicates that the expression level of the IG is found disrupted in the linked cancer type. Upper nodes are color-sorted by their cancer specificity: genes deregulated in a single cancer are shown at the left in blue, followed by genes deregulated in two cancers, tree cancers, and so on. At the right of the upper nodes, there is a single gene (HASPIN) whose expression is disrupted in all tissues.

**Supplementary Figure 3.**
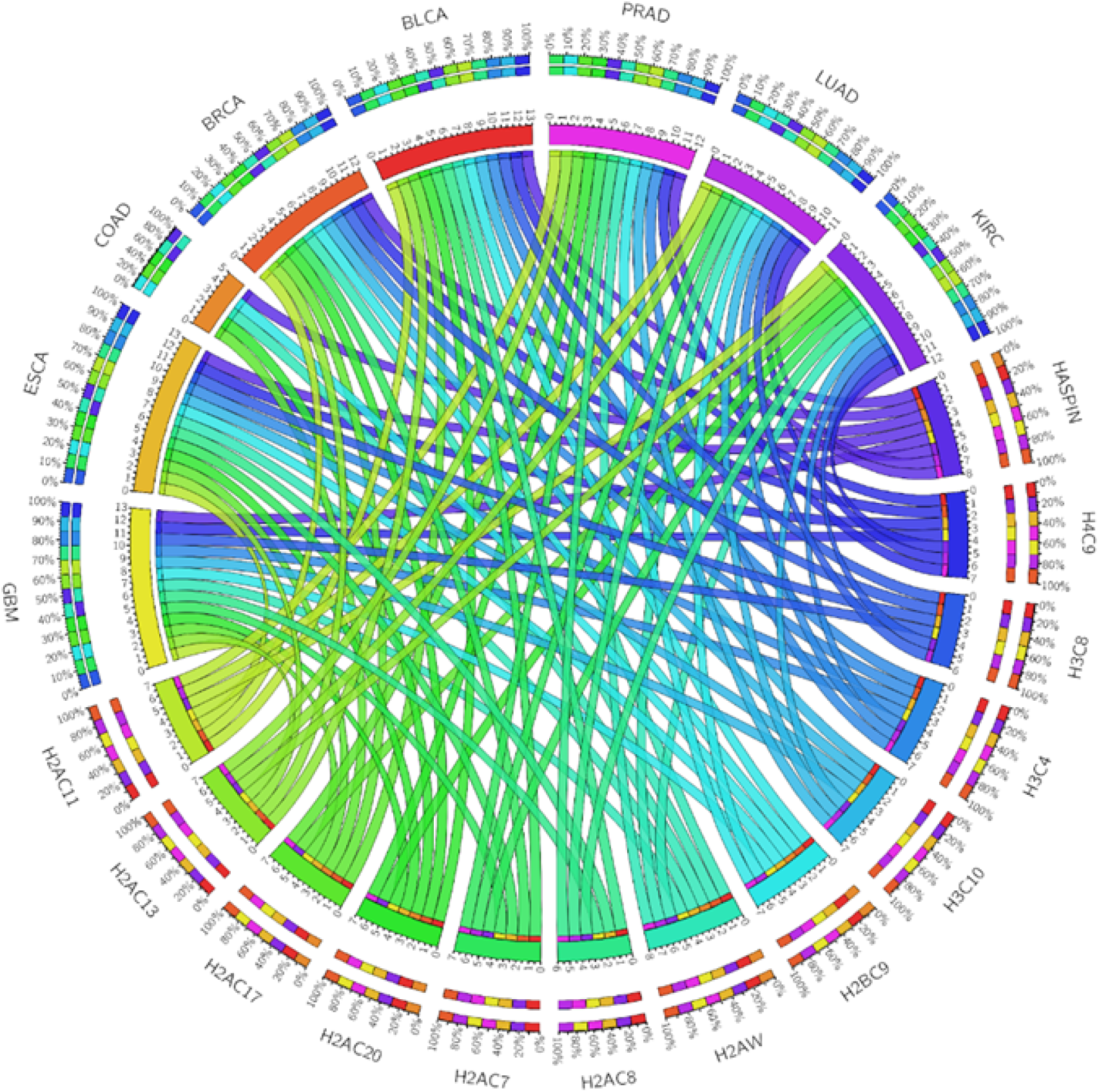
A high conserved group of up-regulated histones across tumors. Circa plot depicts the conservation of the deregulation of a group of histones across cancers. In the upper hemisphere, the total number of histone-related DE-IGs in each cancer is represented by an arc: GBM (yellow), ESCA (mustard), COAD (orange), BRCA (dark orange), BLCA (red), PRAD (pink), LUAD (violet), and KIRC (purple). Histone encoded IGs are depicted in the lower hemisphere. The inner arcs represent the distribution of the histones in each cancer (0-13 genes), while the outside arcs represent the tumors that have that particular gene upregulated (on a scale of 0-100%). Data was obtained from the TCGA database

**Supplementary Figure 4.**
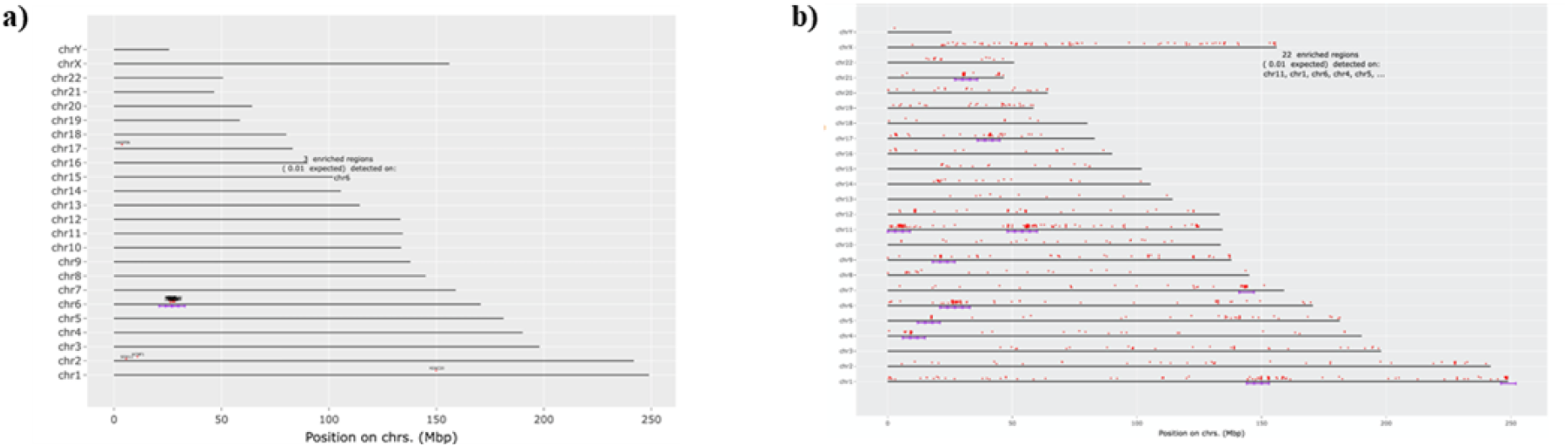
Comparison of genome location of highly conserved DE-IGs. a) Genome location of IGs in the human genome. b) Genome location of highly conserved DE-IGs IGs encoded proteins are represented by red dots. The purple lines indicate regions where these genes are statistically enriched, compared to the density of genes in the background. The hypergeometric test is used to determine if the presence of the genes is significant. Essentially, the genes in each region define a gene set/pathway. The chromosomes may be only partly shown as the last gene’s location to draw the line is used. Data was obtained from the ENSEMBL database.

**Supplementary Figure 5.**
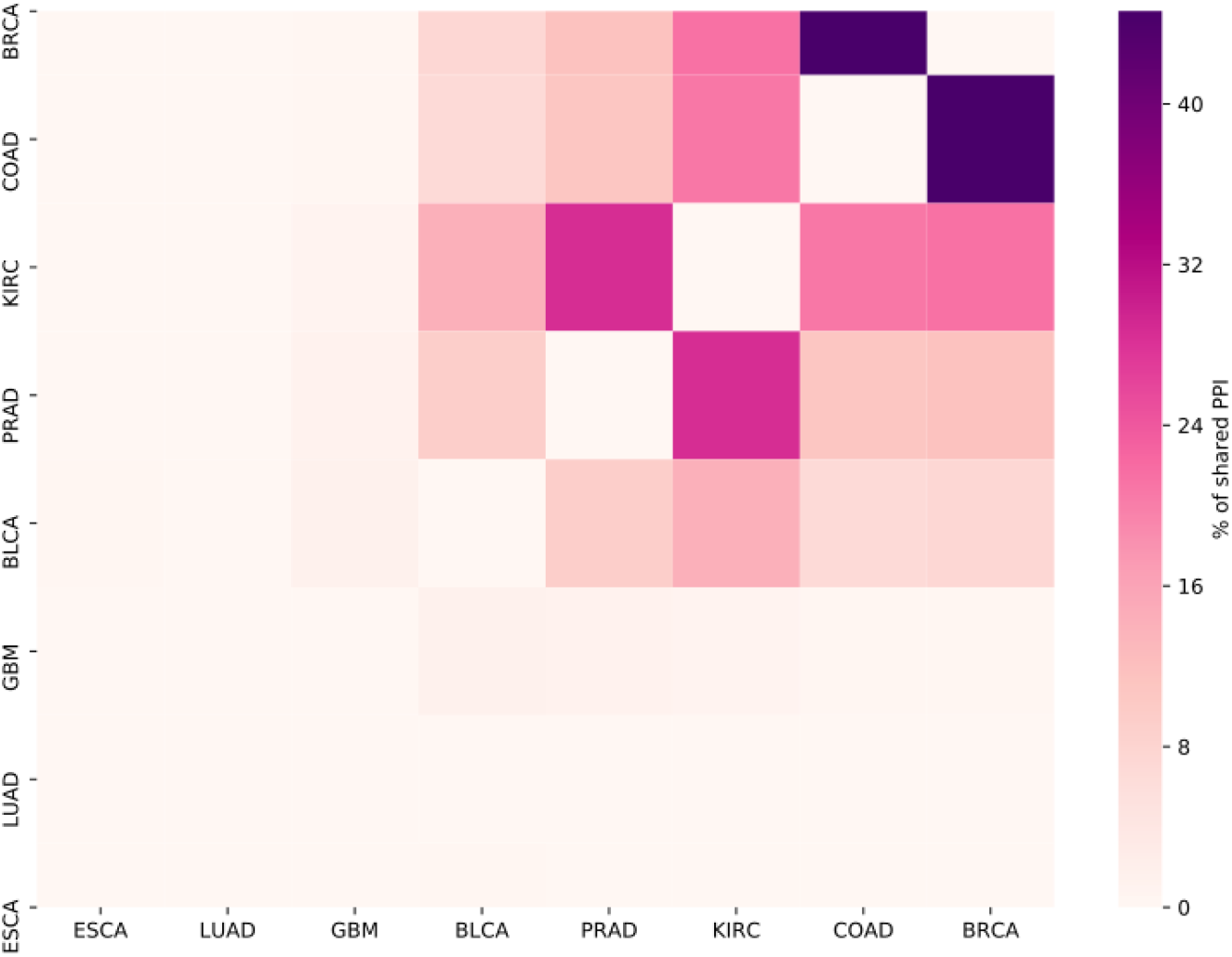
Shared PPI heatmap. Color intensity indicates a higher percentage of shared PPI related to DE IGs of the compared diseases. LUAD does not share any DE IG interaction with other cancer types, and ESCA only one with BCLA. GMB shares almost no interaction with any other cancer (a maximum of 1.64 % with BLCA). COAD and BRCA are the two cancers where more DE IGs are shared, therefore having more interactions in common.

**Supplementary figure 6.**
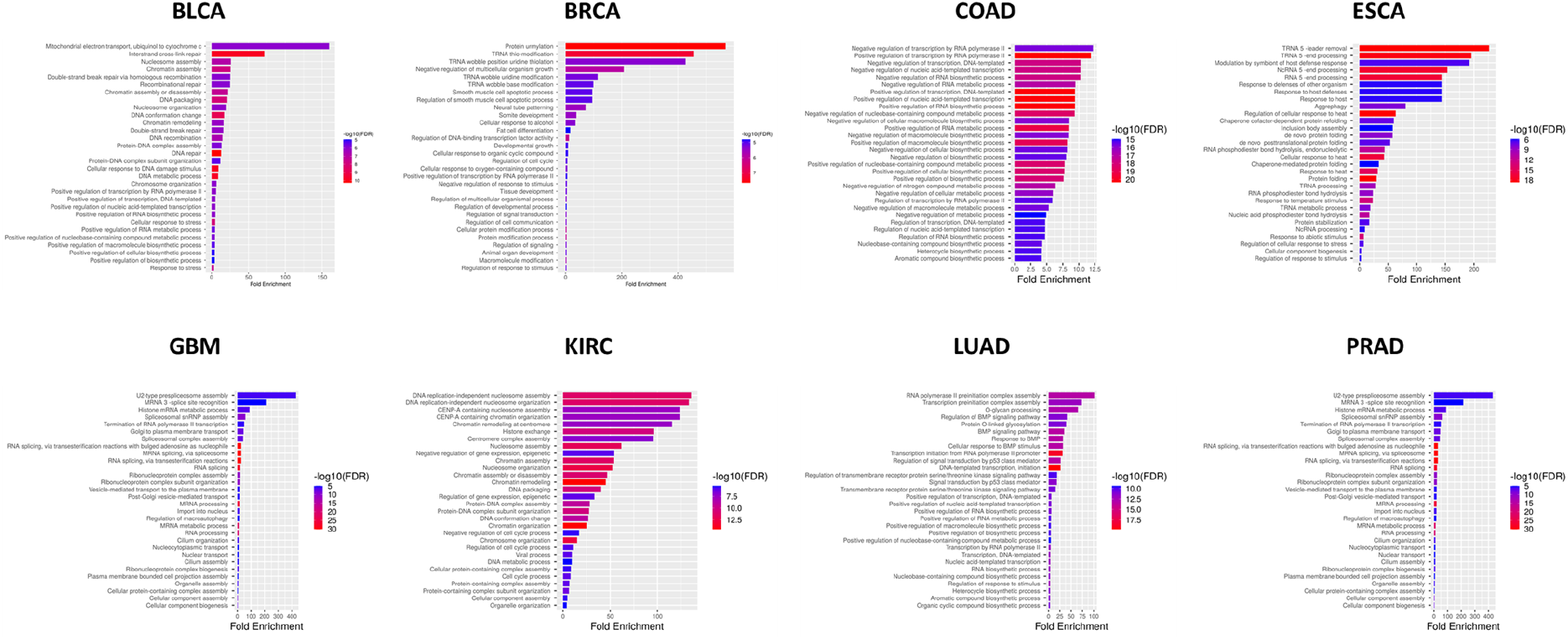
PPI networks ontologies. FDR is calculated based on nominal P-value from the hypergeometric test. Fold Enrichment is defined as the percentage of genes in each cancer belonging to a pathway, divided by the corresponding percentage in the background. FDR indicats how likely the enrichment is by chance. In x-axis the Fold Enrichment indicates how drastically genes of a certain pathway are overrepresented. FDR values are depicted red (high) to blue (low).

**Supplementary Table 1.** Available expression data for each type of cancer.

| Cancer Type | Primary Solid Tumor Samples | Solid Tissue Normal Samples |
| --- | --- | --- |
| BLCA | 414 | 19 |
| BRCA | 1102 | 113 |
| COAD | 478 | 41 |
| ESCA | 161 | 11 |
| GBM | 156 | 5 |
| KIRC | 538 | 72 |
| LUAD | 533 | 59 |
| PRAD | 498 | 52 |

**Supplementary table 2.**
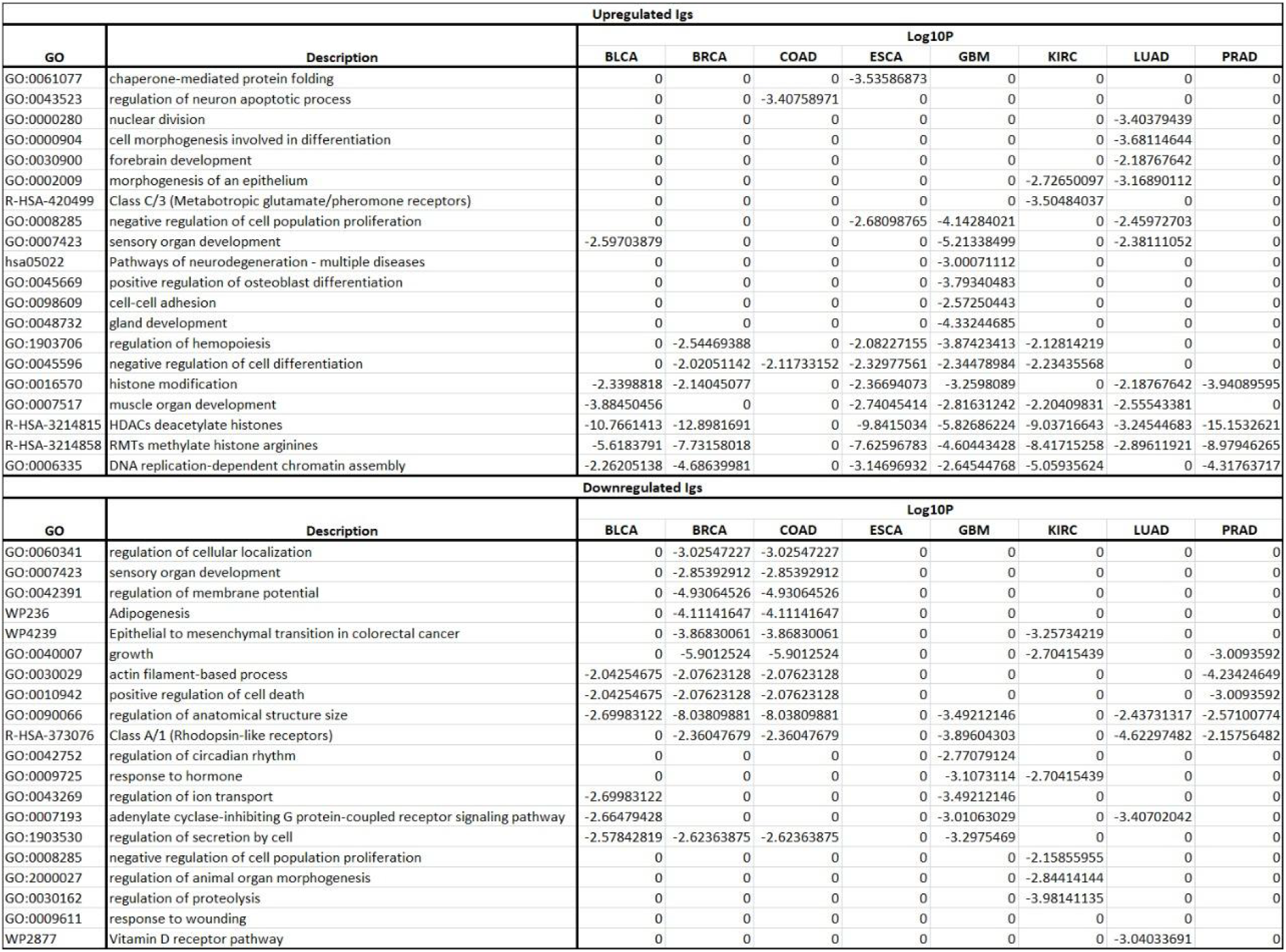
GO enrichment of deregulated IGs.

**Supplementary Table 3.**
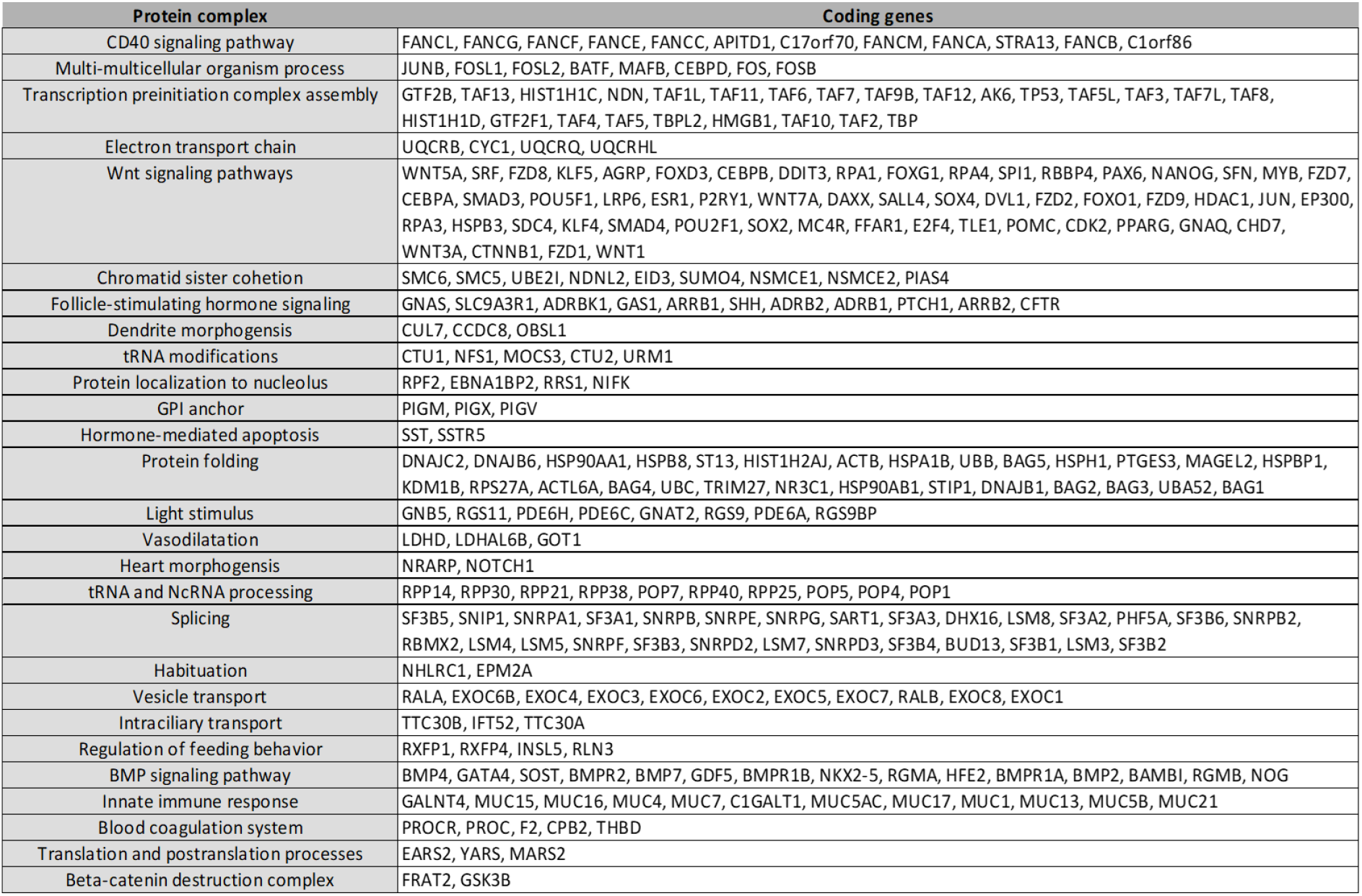
Cancer-specific deregulated genes. Each column contains the list of cancer-specific DE intronless genes.

**Supplementary Table 4.** Shared PPIs. The first column contains pairs of cancers, followed by the number of PPI related to unique DE IG of the compared diseases, finally, in the third column is shown the percentage of shared interactions. Note that two cancer may share more genes than other couples of genes but have a smaller percentage, this is due to a difference in the total number of interactions associated with the compared tumors, for example, PRAD and KIRK share 227 PPIs, representing a percentage of 28.64, while COAD and BRCA have less common interactions (145) but a bigger percentage: 44.61.

| cancers | Interactions shared | Percentage of total interactions in compared cancers |
| --- | --- | --- |
| COAD,BRCA | 145 | 44.6153846153846 |
| KIRC,PRAD | 277 | 28.6452947259566 |
| KIRC,BRCA | 132 | 21.3247172859451 |
| KIRC,COAD | 138 | 20.7518796992481 |
| BLCA,KIRC | 118 | 14.1656662665066 |
| BRCA,PRAD | 95 | 11.6421568627451 |
| COAD,PRAD | 95 | 10.9447004608295 |
| BLCA,PRAD | 95 | 9.3503937007874 |
| BLCA,BRCA | 43 | 7.47826086956522 |
| BLCA,COAD | 43 | 6.85805422647528 |
| BLCA,GBM | 14 | 1.64512338425382 |
| PRAD,GBM | 15 | 1.31233595800525 |
| KIRC,GBM | 9 | 0.910010111223458 |
| BRCA,GBM | 2 | 0.301659125188537 |
| COAD,GBM | 2 | 0.27972027972028 |
| BLCA,ESCA | 1 | 0.175131348511383 |

**Supplementary Table 5.**
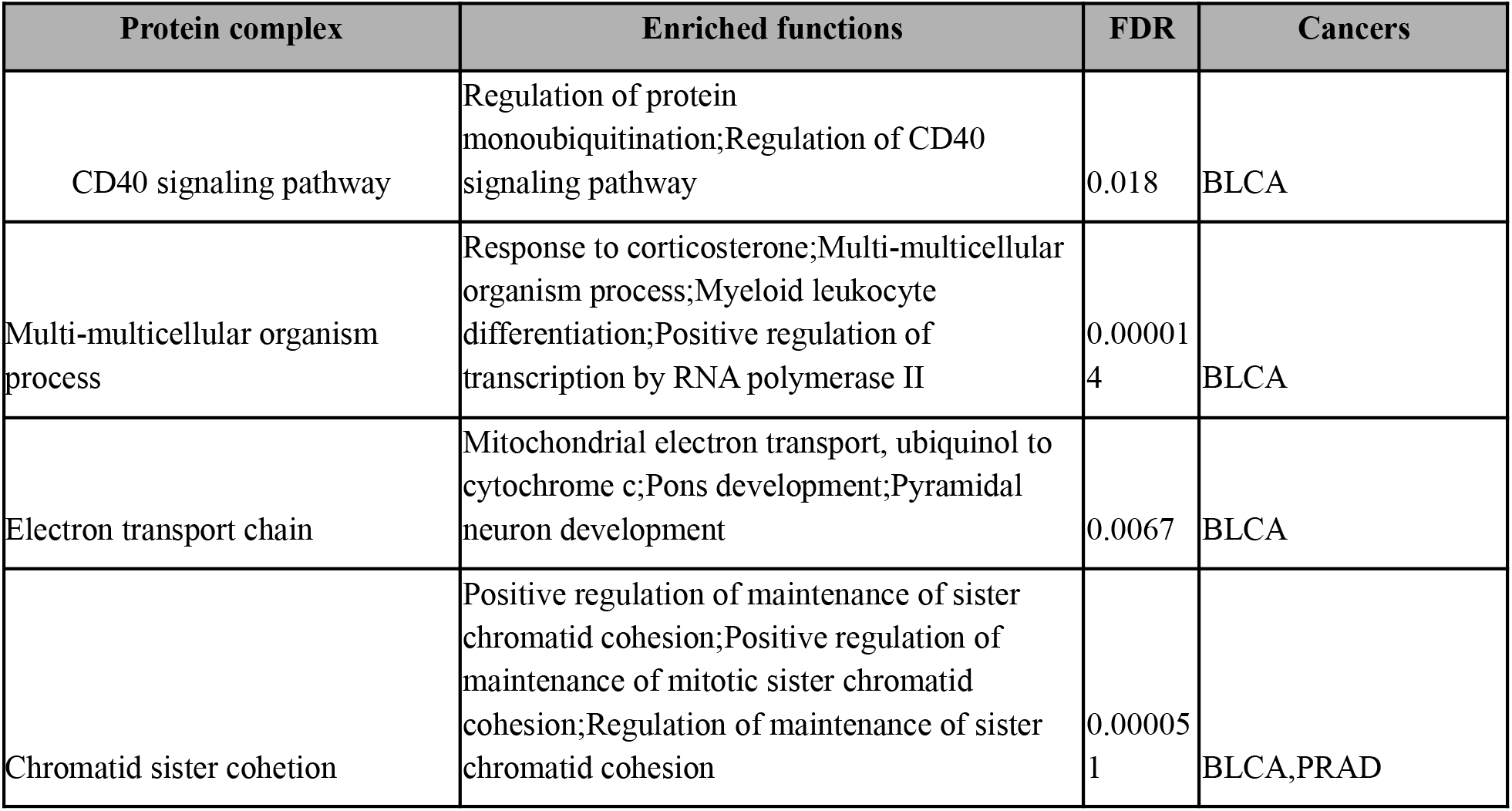
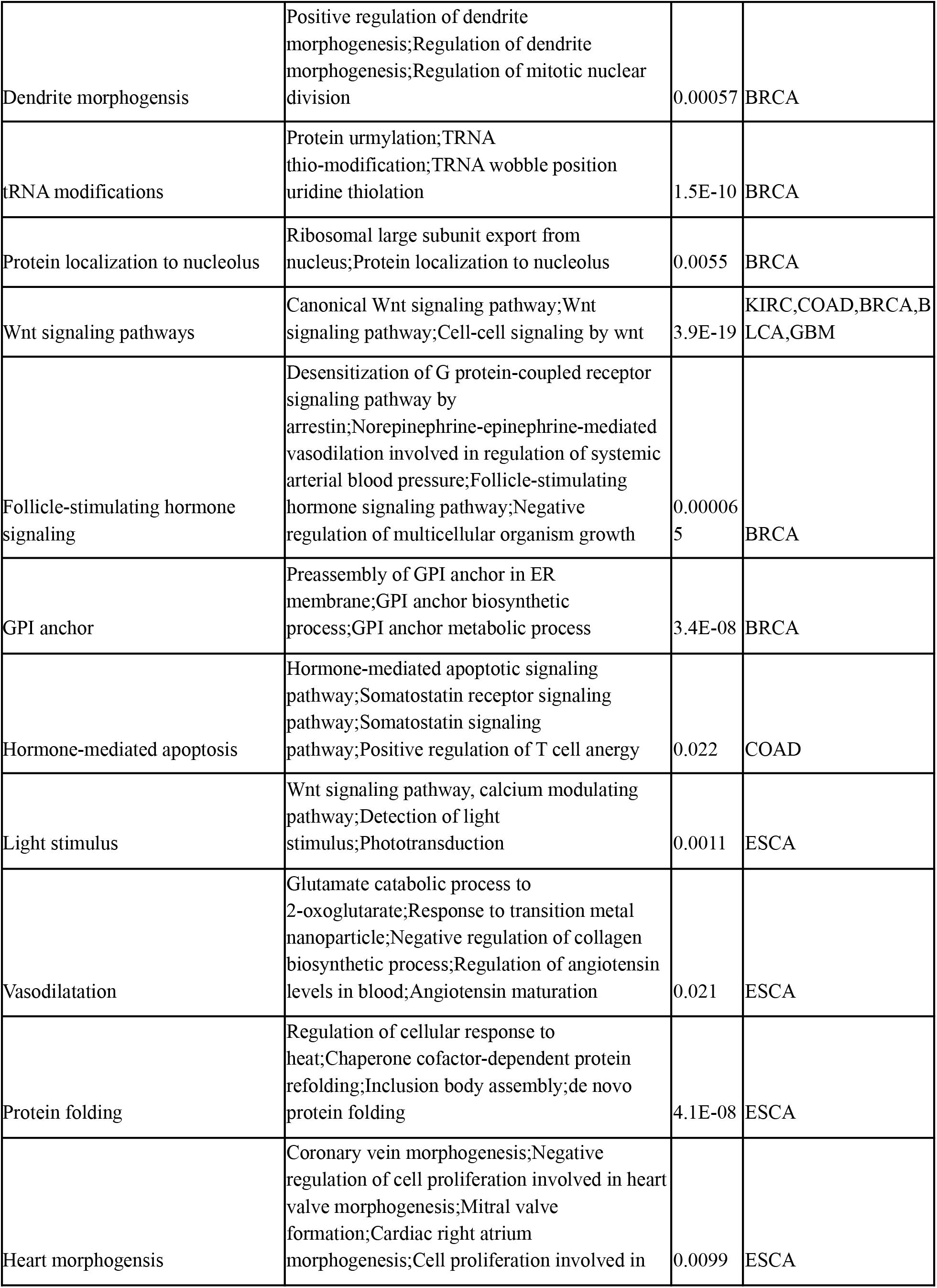
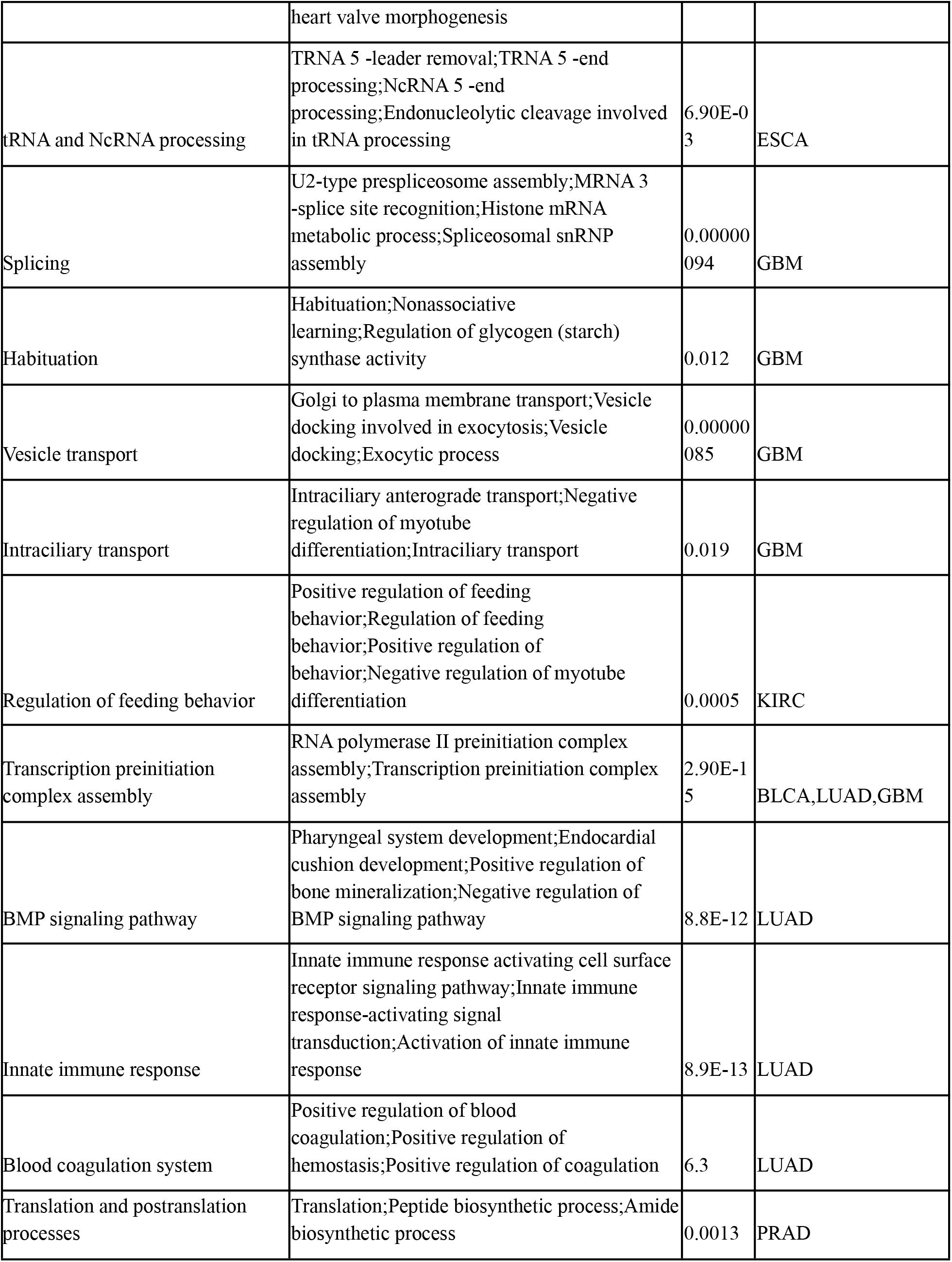
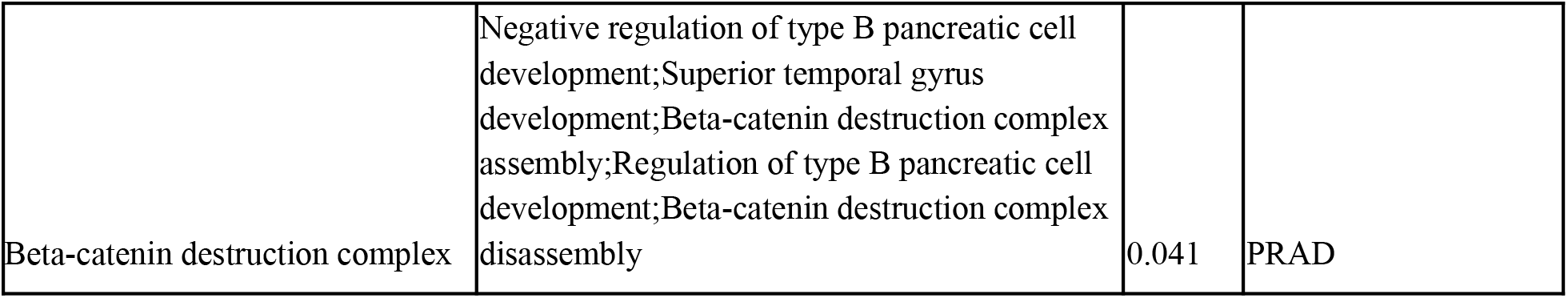
Identified PPI complexes. Protein complexes identified where cancer-specific IG-encoded proteins and their interactors play a key role.

